# Cellular compartment analysis of temporal activity by fluorescent in *situ* hybridization (catFISH) in the transcardially perfused rat brain

**DOI:** 10.1101/767111

**Authors:** Ali Gheidi, Vivek Kumar, Christopher J Fitzpatrick, Rachel L Atkinson, Jonathan D Morrow

**Author notes:** Corresponding Author Jonathan D Morrow, Biomedical Science Research Building, room 5047, 109 Zina Pitcher Place, Ann Arbor, MI 48109. Declarations of interest: none.

## Abstract

Cellular compartment analysis of temporal activity by fluorescent in *situ* hybridization (catFISH) allows high spatiotemporal resolution mapping of immediate early genes in the brain in response to internal/external stimuli. One caveat of this technique and indeed other methods of *in situ* hybridization is the necessity of flash-freezing the brain prior to staining. Often however, the mammalian brain is transcardially perfused to use the brain tissue for immunohistochemistry, the most widely-used technique to study gene expression. The present study illustrates how the original catFISH protocol can be modified for use in adult rats that have been transcardially perfused with 4% paraformaldehyde. c-Fos activity induced by either an auditory tone or status epilepticus was visualized using the catFISH procedure. Analysis of the rat prefrontal cortex, hippocampus and amygdala shows that a clear distinction can be made between the compartmental distribution of c-Fos mRNA in the nuclei and cytoplasmic regions. Furthermore, the qualitative proportion of c-Fos compartmentalization is similar to previous reports of c-Fos expression pattern in rodents navigating novel environments. c-Fos catFISH on perfused rodent brains is an valuable addition to the traditional histological methods using fluorescently labeled riboprobes, and opens several avenues for future investigations.

## 1.0 Introduction

*In situ* hybridization (ISH) has been extensively used to visualize the expression of immediate early genes (IEGs) like c-Fos, c-Jun and activity-regulated cytoskeleton-associated protein (Arc) across brain regions, providing a means to map neural activity in response to a variety of external stimuli and during complex behaviors. Sensory stimuli induce c-Fos in the central nervous system (Hunt, Pini, & Evan, 1987), and c-Fos is a critical regulator of activity-dependent transcriptional programs, synaptic plasticity, and associative learning (Cohen & Greenberg, 2008; Herdegen & Leah, 1998; Morgan & Curran, 1989). The unique ability of IEGs to provide large-scale functional mapping of neural activity at a cellular resolution level has been exploited in numerous neuroscientific studies. Despite advances in IEG-mapping strategies, such as transgenic reporter lines of c-Fos (Barth, 2007), ISH or immunohistochemistry (IHC) continue to be the most widely-used techniques to adequately detect and report IEG signals. The development of cellular compartment analysis of temporal activity by fluorescence ISH (catFISH) of IEGs has further advanced the imaging of neural activity by adding a temporal dimension to an otherwise static technique. Due to the temporal dynamics of translation and translocation, neurons active shortly before sacrifice express IEG mRNA in the nucleus, while those with IEG mRNA that has translocated to the cytoplasm were active at a prior time point, and neurons that have both intranuclear and cytoplasmic IEG mRNA must have been active at both times. Thus, neural activity in response to stimuli at different time points can be distinguished based on the intracellular distribution of mRNA transcripts (Guzowski, McNaughton, Barnes, & Worley, 1999).

Advancement of the classic techniques described above has been somewhat limited due to the manner with which brain tissue must be prepared for a given technique. For ISH, brains are usually fresh-frozen (not fixed) to preserve the integrity, cellular morphology, and spatial composition of macromolecules, such as mRNA, within tissue specimens (Fox, Johnson, Whiting, & Roller, 1985). In contrast, transcardial perfusion, particularly with paraformaldehyde, has several advantages over flash freezing and has therefore become the standard for most immunohistochemical procedures. For example, flushing the brain with paraformaldehyde removes endogenous peroxidase making blocking easier for antigen detection methods such as DAB (3,3’-diaminobenzidine). Further, homogeneous fixation through perfusion of the whole brain yields more uniform and consistent results across subjects. Performing ISH on perfused brains could enable an effective combination of these powerful techniques, for example by processing adjacent sections for ISH and IHC on the same subject. However, formaldehyde fixation also results in base pair modifications that prevent the base-pairing with riboprobes required for ISH. More specifically, formaldehyde fixation results in cross-linking of nucleic acid bases to other macromolecules (Feldman, 1973) in a two-step reaction involving (1) the formation of a hydroxylmethyl group (N—CH_2_OH) through the addition of a formaldehyde group to the base, and (2) electrophilic addition between the newly formed hydroxylmethyl group and amino groups of other bases resulting in methylene bridges (Auerbach, Moutschen-Dahmen, & Moutschen, 1977)). In addition, mRNA bases are differentially susceptible to these reactions (Adenine, Cytosine ≫ Guanine > Uracil) with hydroxymethylation (and methylene bridge formation) ranging from 4% for uracil to 40% for adenine (Masuda, Ohnishi, Kawamoto, Monden, & Okubo, 1999). Multiple fixatives have been utilized over the decades (i.e., aldehydes, oxidizing agents, and alcohol-based fixatives); however, to date, no fixative has been discovered that preserves the integrity and spatial composition of mRNA without also reducing the detection/sensitivity of riboprobes.

Antigen retrieval protocols have been extensively used for IHC to “unmask” binding sites and allow better penetration of antibodies in perfused samples. Using IHC, antigen retrieval with proteinase K treatment has been demonstrated with c-Fos protein in rat spleen tissue following whole-animal perfusion fixation (Meltzer, Grimm, Greenberg, & Nance, 1997); however, there has not yet been an adaptation of antigen retrieval for ISH using the catFISH method and whole-animal perfusion. Given the fact that IHC and ISH require different tissue processing, including preparation (i.e., free-floating versus slide-mounted) and section thickness, optimizing antigen retrieval concomitantly for each technique will be crucial for moving the field forward. The protocol described here allows for mRNA measurement in perfused brain tissue with sufficient resolution to perform catFISH.

## 2.0 Materials and Methods

### 2.1 Animals

Adult male Sprague Dawley rats (275-300 g) were purchased from Charles River Laboratories. Rats (two per cage) were maintained on a 12 h light/dark cycle, and standard rodent chow and water were available *ad libitum*. All procedures were approved by the University Committee on the Use and Care of Animals (University of Michigan; Ann Arbor, MI).

### 2.2 Seizure Induction

We used a seizure model, induced pharmacologically as described previously (Althaus, Zhang, & Parent, 2016; Kron, Zhang, & Parent, 2010) to perform a qualitative comparison between our current modification and previous catFISH protocols. Briefly, rats were pretreated with atropine methylbromide (5 mg/kg, i.p.; Sigma-Aldrich, Inc.; St. Louis, MO) 20 min before seizure induction with pilocarpine hydrochloride (345 mg/kg, i.p.; Sigma-Aldrich, Inc.).

### 2.3 Tone-Induced c-Fos Expression

To evaluate the suitability of our current modification for quantification purposes, we exposed a group of rats to a 6-minute tone in an operant box, then returned the rats back into the home cage and sacrificed 30 minutes later. The results from this group were quantified and used for statistical analyses.

### 2.4 Tissue Preparation

Rats were deeply anesthetized with a solution (1 mL/kg; i.p.) of ketamine (90 mg/kg) and xylazine (10 mg/kg), and euthanized by transcardial perfusion with saline immediately followed by a solution of 4% paraformaldehyde (PFA) in 0.1 M phosphate-buffered saline (PBS; pH = 7.34-7.36). Brains were extracted from the skull and post-fixed in 4% PFA in 0.1 M PBS overnight. Next, brains were saturated in a 20% sucrose solution in 0.1 M PBS, flash-frozen in isopentane over dry ice (−30°C), then stored in a −80°C freezer until further processing. Brains were sectioned using a cryostat (CM1860; Leica Biosystems; Buffalo Grove, IL) at 20 µm and thaw-mounted on Fisherbrand™ Superfrost™ Plus slides (Fisher Scientific; Waltham, MA).

### 2.5 Preparation of c-Fos Riboprobes

*In vitro* transcription was performed as previously described (Guzowski & Worley, 2001).

#### Day 1

##### Prehybridization Steps

\# 20-µm cryosectioned^1^ slides should be brought from −80°C to −20°C prior to thawing at room temperature for ten minutes.

\# Steps on Day 1 which should be performed with DEPC H_2_O^2^ or molecular grade H_2_O are indicated as [DEPC H_2_O]

1. Place slides in proteinase K^3^ (final concentration 500 µg/50mL of proteinase k buffer)^4^ at 37°C for 30 min
2. 5 minutes in proteinase K buffer at room temperature
3. 5 minutes in 4% paraformaldehyde^5^ at 4°C [DEPC H_2_O]
4. 2 minutes in 2X SSC^6^ at room temperature [DEPC H_2_O]
5. 10 minutes in 0.5% acetic anhydride^7^ at room temperature [DEPC H_2_O]
6. Briefly rinse in DEPC H_2_O
7. 5 minutes in acetone: methanol^8^
8. 5 minutes in 2X SSC [DEPC H_2_O]

##### Hybridization Steps

9. Snap freeze (7 min at 90°C and 5 min on 0°C) ∼300ng^9^ of c-Fos cRNA probes diluted in hybridization buffer^10^
10. Apply the denatured probe solution on to the slides and incubate^11^ overnight at 56°C

#### Day 2

Use a shaker for all room temperature washes

1. Bring slides to room temperature
2. Remove coverslips and wash 4X with 2XSSC for 15 minutes each
3. Wash in 2X SSC and RNase A^12^ for 30 min at 56°C
4. Wash in 2X SSC for 10 minutes at room temperature
5. Wash in 0.5X SSC for 10 minutes at room temperature
6. Wash with 0.5X SSC for 30 minutes at 70°C
7. Wash with 0.5X SSC for 5 minutes at room temperature
8. Quench endogenous peroxidase with 2% H_2_O_2_ ^13^ for 15 minutes at room temperature
9. Wash 2 X with 1X SSC-Tween^14^ at room temperature for 5 minutes
10. Wash 5 minutes with 0.1M TBS^15^
11. Draw hydrophobic barriers around brain sections using a PAP ^16^pen
12. Block brain tissue sections with 0.5% TSA blocking^17^ buffer and 5% normal sheep serum for 30 minutes at room temperature
13. Apply antibody (1:400)^18^ and incubate at room temperature for 2 hours or overnight at 4°C
14. Remove coverslips and wash 4X with 0.1M TBS-T (tween 0.05%)^19^ for 15 minutes at room temperature
15. Apply TSA Cy3^20^ system for 30 minutes at room temperature
16. Wash 4 X with 0.1M TBS-T for 15 minutes at room temperature
17. Wash 1 X with 0.1M TBS for 5 minutes at room temperature
18. Counterstain with DAPI^21^ for 30 minutes at room temperature
19. Wash 1X with 0.1M TBS for 10 minutes at room temperature
20. Apply anti-fade mounting media^22^ on slides and seal edges with clear nail polish
21. Slides are ready for imaging. Keep slides in dark at 4°C when not imaging.

#### Appendix

##### 0.5% Acetic Anhydride

Make final volume of 0.92% NaCl^23^, 1.5% Triethanolamine^24^ and add acetic anhydride^25^ at 0.5% just prior to use

##### 20X SSC

Make final volume with 17.52% NaCl^21^ and 8.82% sodium citrate^26^

##### 4% paraformaldehyde

Heat water to 56°C and add 4% paraformaldehyde powder^27^. Add 0.44% NaOH^28^, <u>wait to clear</u> and add 1.6% NaH_2_PO_4_

##### Protease K buffer

Use one to one: Dilute 1M Tris HCl to 50mM with water. Dilute 0.5M EDTA^29^ with water to 5mM (keep separate until used for procedure). Make stock aliquots of 1mg/mL of proteinase K^2^ in both buffers combined. Add proteinase K^2^ to 50mL of both buffers combined for working solution

### 2.6 Image Acquisition and Cell Counting

Images were acquired on a Zeiss® AxioImage M2 fluorescent microscope with an Apotome® using fluorescent filters for cubes for DAPI (or Alexa 350, AMCA, BFP, Hoechst,) shift free with band pass emission filter and Cy3 (c-Fos) and using a 10X/ 20X or 40X objective. Images were processed and analyzed with ImageJ software (NIH). DAPI positive cells were segmented into binary form and counted automatically using the ‘analyze particle’ plugin of ImageJ (See supplemental content). An experimenter blinded to the conditions performed manual counting of c-Fos as either-1) nuclear, 2) cytoplasmic, or 3) both nuclear and cytoplasmic (Dual) (Marrone, Schaner, McNaughton, Worley, & Barnes, 2008).

### 2.7 Statistical Analysis

‘IBM® SPSS® 24’ software was used for the statistical analysis of compartmental mRNA expression. Within-subject comparisons were made as previously described (Marrone et al., 2008). Briefly, comparisons of 1) c-Fos positive cells in the nuclei, 2) cytoplasm 3) both nuclei and cytoplasm [All as a fraction of total cells counted] were made using paired sample t-tests. A similar test was performed to compare the total active cells in the home cage (c-Fos positive in nuclei, and dual nuclei and cytoplasmic) to cells active during tone (c-Fos positive in cytoplasm, and dual nuclei and cytoplasmic).

To examine the differences, paired sample t-tests were run between the percentage of c-Fos in each compartmental region (e.g., intranuclear *vs* cytoplasmic) and also the amount of total active cells during tone and home cage. No between-subjects factor was used here. Significance level was set at *p<0.05* for detecting effects.

## 3. Results

c-Fos expression was clearly evident in rats given status epilepticus (Figure 1 A). In contrast, perfused brains subjected to previous catFISH protocols failed to generate a clear Cy3 signal to detect c-Fos levels (Figure 1B). To verify whether patterns of c-Fos distribution as seen using our protocol matches catFISH results from previous reports, we exposed rats to an auditory tone and placed them back in their home cages for 25 min before sacrifice. This provided with a more realistic spatiotemporal dimension that we cannot get with a seizure protocol. Using this modified protocol, we were able to detect IEG expression in the hippocampus, amygdala and prefrontal cortex (Figure 2). We found that the qualitative distribution of c-Fos closely resembled the distribution previously reported with another IEG, Arc (Guzowski et al., 1999).

**Figure 1.**
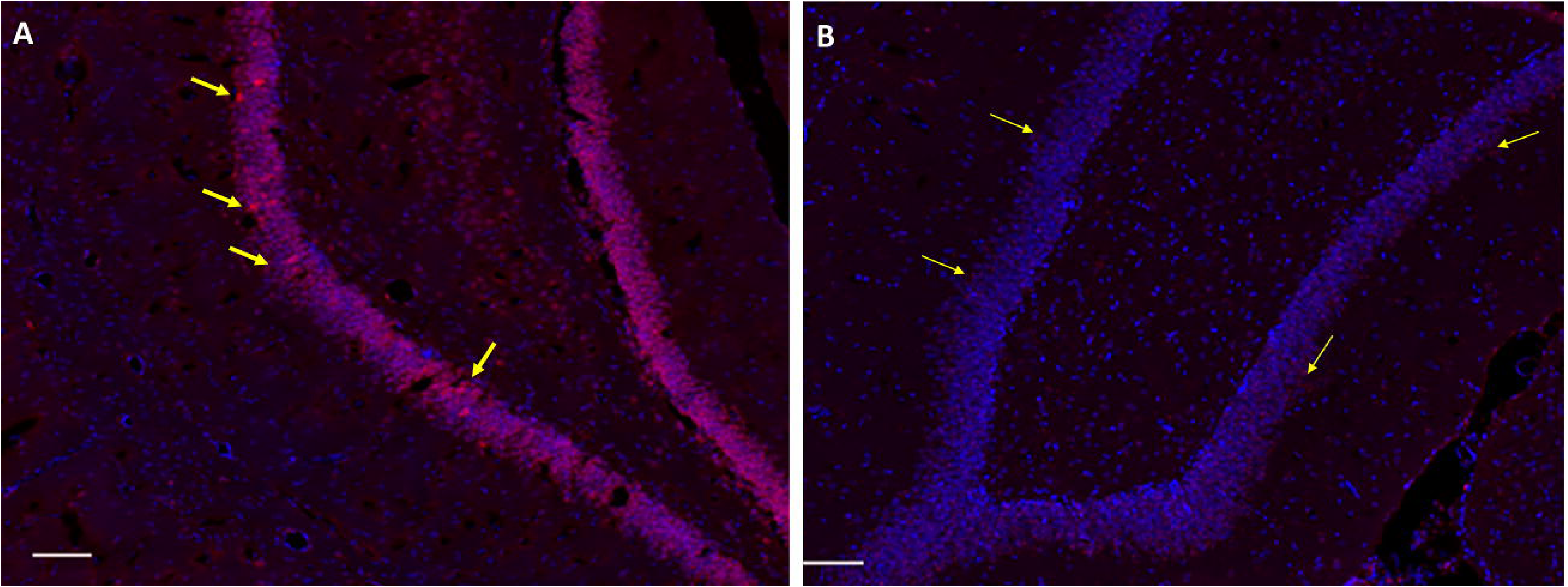
c-Fos catFISH in perfused dentate gyrus of rat following 25 minutes of seizure. Figure 1A (thick arrows) shows brighter red signal of c-Fos in dentate gyrus following the current protocol as opposed to weaker red signal (Figure 1B, thin arrows) following Worley & Guzowski, 2001 protocol. Blue=DAPI, Red=Cy3. Scale bar-100 µm.

**Figure 2.**
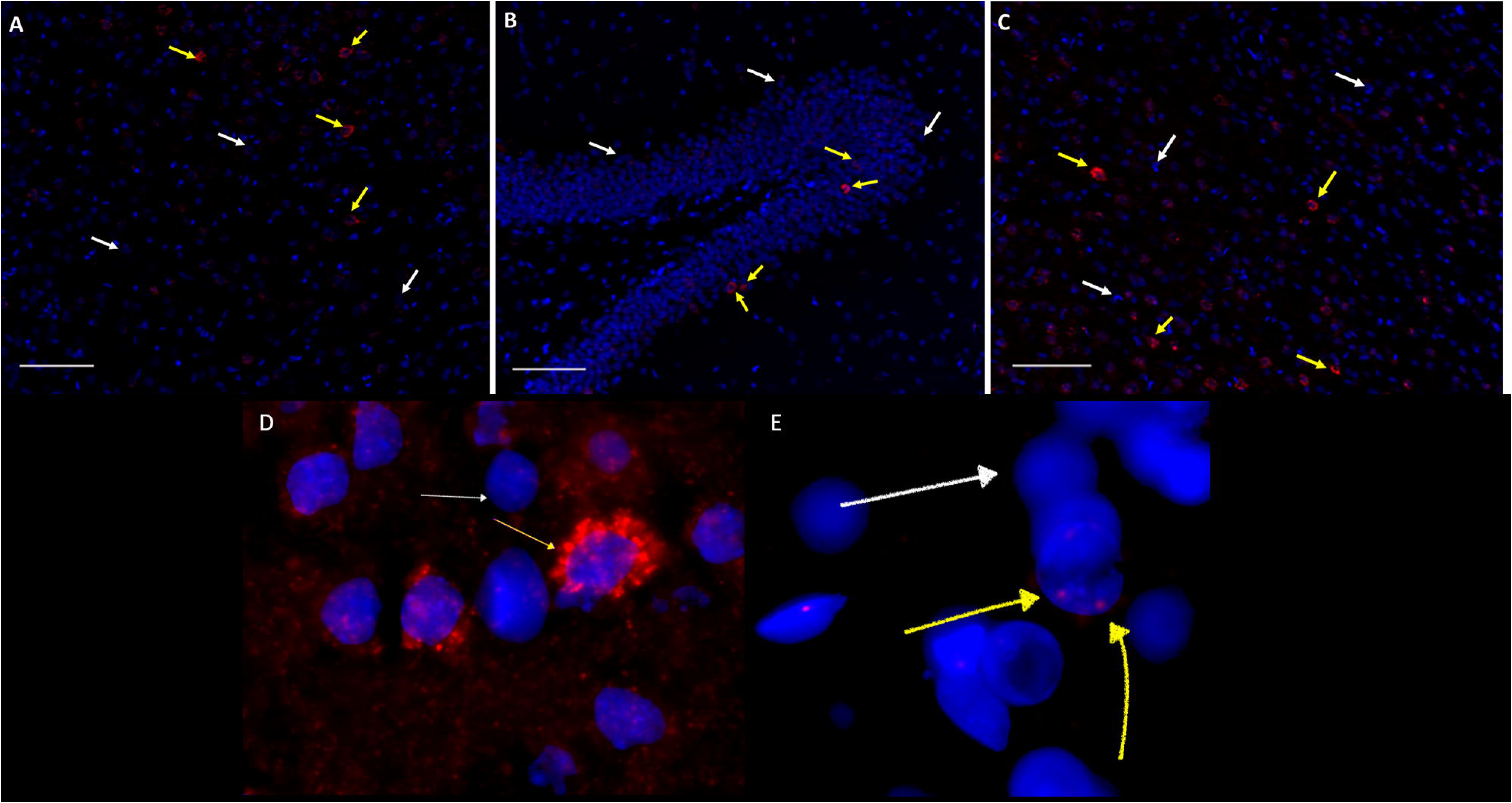
c-Fos activity in perfused rat brains given an auditory tone and sacrificed 30 minutes later. Images from the amygdala (A), hippocampus (B) and prefrontal cortex (C) show discernable signal between c-Fos positive (red signal, yellow arrows) and c-Fos negative (blue signal, white arrows). Bottom panels show high-magnification (40X) examples of a cell containing only cytoplasmic (D) or only nuclear (E) staining. Blue=DAPI, Red=Cy3. Scale bar-100 µm

As expected, paired sample t-tests showed that the percentage of cells with c-Fos expression in the cytoplasmic compartment (1.65, SD=1.38) was higher than the percentage of intra-nuclear (M=0.69, SD=0.57), t (24) = −3.8, p =0.001 and double c-Fos-positive cells (M=1.05, SD=1.18) t (24) =3.45, p=0.002 (Figure 3). Because the mRNA expression in cytoplasmic compartment is thought to correspond to the time of tone delivery 30 min prior to sacrifice, these results indicate that quantitative compartmental expression of c-Fos was reflective of sensory experiences prior to death. Our results are also consistent with previous work in rats given a different behavioral paradigm (ambulatory exploration of an environment) and sacrificed 30 minutes later, showing that the proportion of c-Fos was highest in the cytoplasmic compartment (Gheidi, Satvat, & Marrone, 2012; Marrone et al., 2008). Further, we calculated the total number of c-Fos positive cells in the cingulate cortex that were active during tone (Cytoplasmic + Double) (M=2.6, SD=2.44) and home cage exposure (Intra-Nuclear + Double) (M=1.73, SD=0.14). We observed significantly higher numbers of cells (t (24) =3.82, p=0.001) that were active in response to the tone as compared to the home cage condition (Figure 4).

**Figure 3.**
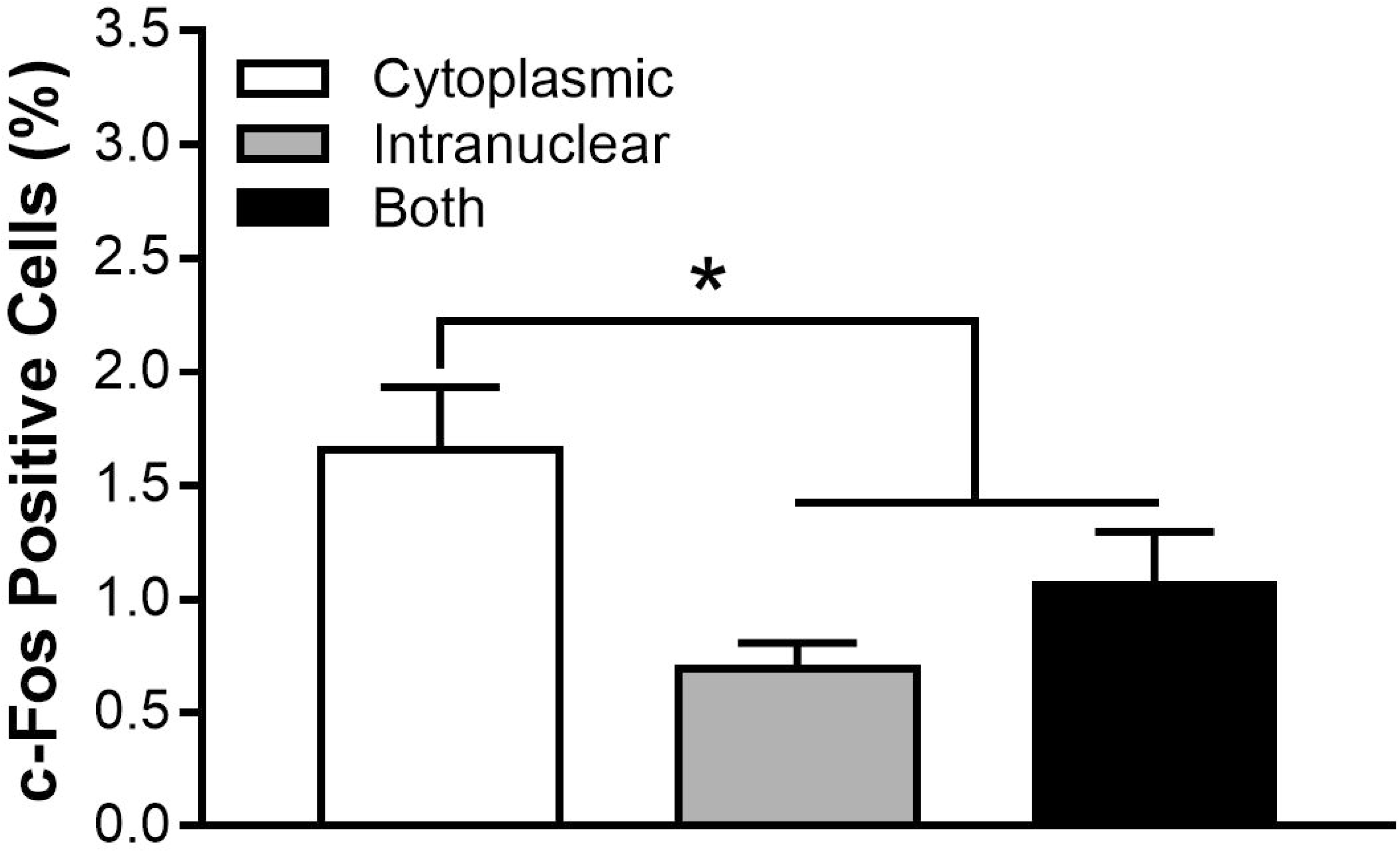
Compartmental distribution of c-Fos mRNA in prefrontal cortex. Rats sacrificed after tone exposure prior to home cage, have higher number of cytoplasmic c-Fos mRNA positive cells. A significant difference between % cytoplasmic mRNA (M=1.65, SD=1.38), and % intra-nuclear (M=0.69, SD=0.57) and % both intra-nuclear and cytoplasmic c-Fos mRNA (M=1.05, SD=1.18) was detected using a paired sample t-test.

**Figure 4.**
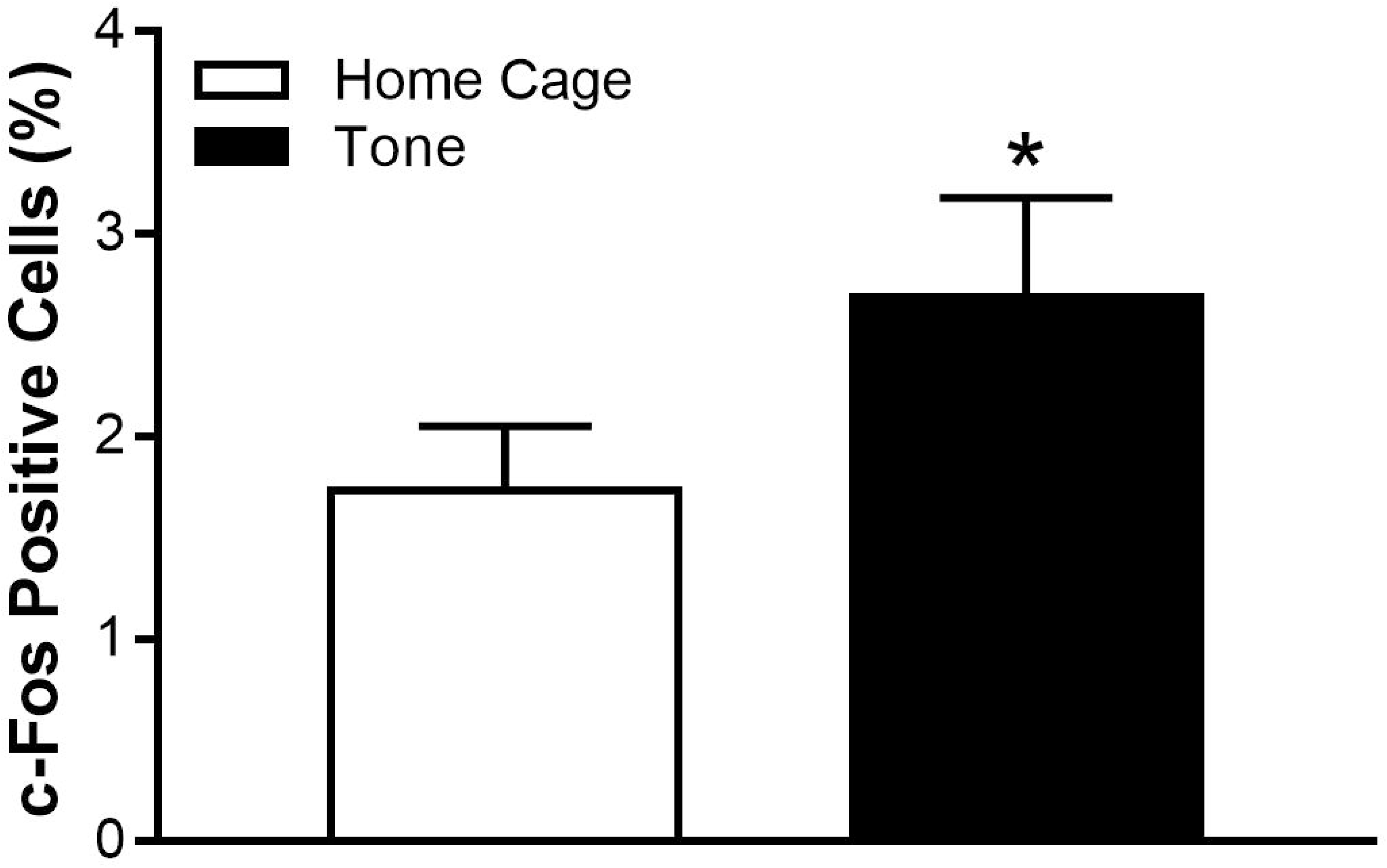
Number of c-Fos positive cells corresponded to behavior prior to sacrifice. Tone exposure prior to rest in the home cage activates expression of c-Fos mRNA. There was a significant difference between cytoplasmic (Tone) (M=2.70, SD= 2.43.) and intra-nuclear (Home Cage) c-Fos positive cells (M=1.73, SD=1.60), as determined by a paired sample t-test.

Lastly, we were interested in determining whether c-Fos activity in response to the tone meant cells were more likely to also be active in the subsequent home cage condition. To test this, we calculated the product of the proportion of cells activated to home cage and tone delivery as an index of the overlap expected due to random chance. This product was then compared to the observed number of cells active in response to both tone and home cage (double labeled cells) using a paired sample t-test. Results showed a significant difference t (24) =-4.69, p=0.000091, between overlap expected by random chance and those observed (double labeled cells). The activity of cells in response to the home cage therefore appears to be related to their prior response to the tone (Figure 5), and not simply due to activation of a random cohort of cells in the home cage. These findings support previous work with both with *in vivo* electrophysiology in freely-moving rodents and IEG studies (Marrone et al., 2008; Wilson & McNaughton, 1994).

**Figure 5.**
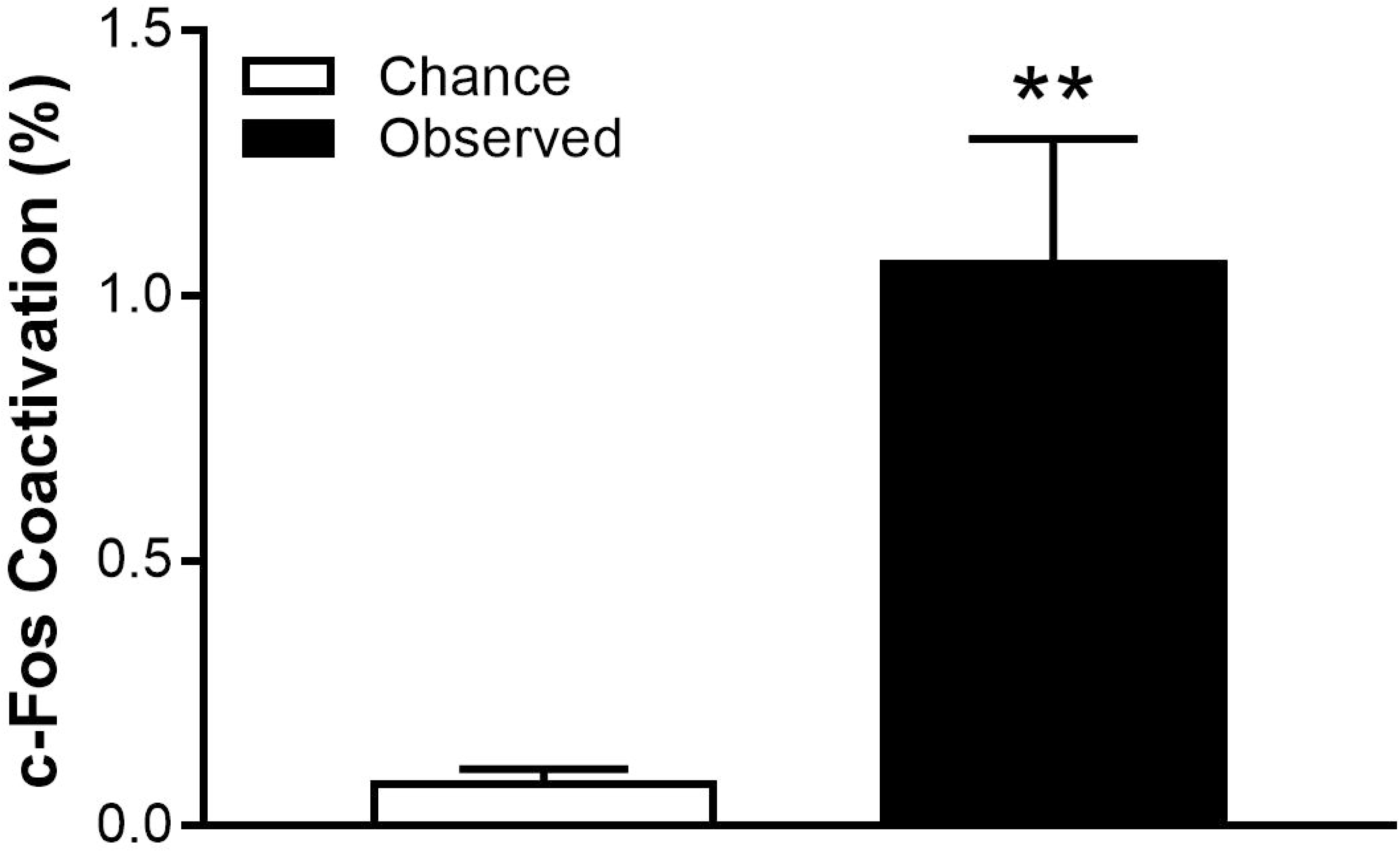
Percent of c-Fos positive neurons activated both during tone and home-cage are far greater than that expected by random chance with replacement. The proportion of cells activated to the tone was multiplied by the proportion of cells activated home cage to get a value expected by chance (M=0.07, SD=0.14) (Marrone et al., 2008). This value was then compared to the number of cells with both intra-nuclear and cytoplasmic mRNA (Double) (M=1.06, SD=1.17) [Observed].

## 4. Discussion

The protocol described here allows for the quantification of c-Fos mRNA in perfused brain tissue with sufficient spatial resolution to perform catFISH. The c-Fos catFISH technique has several advantages over traditional IEG detection methods. For example, relatively time-limited expression of mRNA foci allows tighter temporal resolution of stimuli than protein staining techniques, while the compartmental distribution of c-Fos mRNA allows for within-subject comparisons of responses to different stimuli (Guzowski, McNaughton, Barnes, & Worley, 2001). This spatiotemporal dimension can be used in any experiment in which discrete stimuli can be delivered to the rodent separated by a delay. We and others have used the catFISH technique to track cell assemblies in response to spatial, olfactory and here auditory stimuli. The protocol described above is modified from the (Guzowski & Worley, 2001) method developed for analysis of Arc and Homer 1a genes in fresh frozen tissue. We incorporated changes necessary to optimize the detection of IEG mRNA in perfused fixed brain tissue. The key modifications were based on the following observations: 1) We find that a higher concentration of riboprobe (∼300ng/slide) must be used for perfused brain tissue catFISH. 2) Proteinase K treatment must be performed prior to tissue fixation. 3) By increasing the number of sodium citrate and Tris washes on day 2, and increasing the temperature for both RNase treatment and 0.5X sodium citrate washes, we are able to achieve higher signal intensity and lower background. 4) By using a PAP Pen (ImmEdge, Vector labs) based hydrophobic barrier or silicon hybridization chambers (Secure-seal, Sigma-Aldrich) around tissue sections for hybridization or incubation steps, we achieved better signals than by using a coverslip-based hybridization approach.

Our current finding of compartmental expression of c-Fos is consistent with expression patterns previously reported using fresh frozen brains (Gheidi et al., 2012; Marrone et al., 2008). Interestingly, our c-Fos mRNA positive cell quantification revealed that the number of cells activated in response to both tone and home cage far exceeded that expected by random chance. This may reflect ‘replay’ of the previous experience with tone, as has been previously reported with both *in vivo* electrophysiology in freely moving rats (Wilson & McNaughton, 1994) and IEG detection methods in animals during spatial navigation (Gheidi, Azzopardi, Adams, & Marrone, 2013). The finding lends support to the notion that “background” IEG expression is not mere random activity and may represent mnemonic replay of previous experiences. In the current study, however, it is not clear whether the prefrontal cortical activity was primarily in response to the tone, the operant behavior box, or a combination of both. Further experiments will be necessary to disentangle these confounds. Taken together, the findings presented here suggest that this modified protocol for catFISH in paraformaldehyde perfused fixed rodent brain tissues is sensitive enough to detect the expression of behaviorally relevant genes such as IEGs.

## Supporting information

Supplemental Content

## Acknowledgments

We would like to thank Dr. Stanley Watson and Jennifer Fitzpatrick for their technical support with setting up the catFISH protocol.

## Footnotes

1 Refer to Guzowski & Worley (1999) for sectioning instructions

2 SIGMA D5758-25ML. Add 0.1% of DEPC to molecular grade water

3 Sigma catalogue# P2308-100mg

4 See appendix for protease K buffer

5 Appendix

6 See appendix for 20X SSC stock solution

7 Appendix

8 Should be made prior with a ratio of 1:1 and kept at −20°C prior to use. (Acetone, FISHER A949-1), Methanol (FISHER A452-1)

9 DIG-conjugated CFos riboprobe can be mass verified with a Spectrophotometer (refer to appendix 2 for hapten labeling of CFos)

10 Mix 1:1 with molecular grade formamide (SIGMA)

11 Slides should be elevated in incubation chamber with 1:1 reagent grade formamide and 2XSSC

12 Final concentration is 500µg/50mL. Use 2XSSC (SIGMA R6513-10mg)

13 Dilute with 1X SSC. 30% H_2_O_2_ (SIGMA 216763)

14 For this and all subsequent Tween washes, add 0.05% tween (FISHER BP337) to final volume of buffer

15 Refer to appendix for 1 M Tris-HCl recipe

16 Research Products International Corporation. Item 195505

17 TSA blocking buffer diluent is 0.1M TBS

18 ROCH Anti-Digoxigenin-POD fab fragments Ref: 11207733910. Mix with 0.5% TSA blocking buffer and 0.1M TBS

19 Sigma P9416

20 TSA Cy3 ^PLUS^ has its own diluent. Use 1:50 for each slide (PERKIN ELMER NEL744001KT)

21 1:1000 in 0.1M TBS

22 Appendix

23 FISHER S271

24 SIGMA T58300

25 SIGMA 320102

26 Fisher Catalogue# s279-3

27 Electron Microscopy Sciences Catalogue # 19202

28 FISHER Catalogue # S318-100

29 AMBION part# AM926G

## References

Althaus, A. L., Zhang, H., & Parent, J. M. (2016). Axonal plasticity of age-defined dentate granule cells in a rat model of mesial temporal lobe epilepsy. Neurobiol Dis, 86, 187–196. doi:10.1016/j.nbd.2015.11.024

Auerbach, C., Moutschen-Dahmen, M., & Moutschen, J. (1977). Genetic and cytogenetical effects of formaldehyde and related compounds. Mutat Res, 39(3-4), 317–361.

Barth, A. L. (2007). Visualizing circuits and systems using transgenic reporters of neural activity. Curr Opin Neurobiol, 17(5), 567–571. doi:10.1016/j.conb.2007.10.003

Cohen, S., & Greenberg, M. E. (2008). Communication between the synapse and the nucleus in neuronal development, plasticity, and disease. Annu Rev Cell Dev Biol, 24, 183–209. doi:10.1146/annurev.cellbio.24.110707.175235

Feldman, M. Y. (1973). Reactions of nucleic acids and nucleoproteins with formaldehyde. Prog Nucleic Acid Res Mol Biol, 13, 1–49.

Fox, C. H., Johnson, F. B., Whiting, J., & Roller, P. P. (1985). Formaldehyde fixation. J Histochem Cytochem, 33(8), 845–853. doi:10.1177/33.8.3894502

Gheidi, A., Azzopardi, E., Adams, A. A., & Marrone, D. F. (2013). Experience-dependent persistent expression of zif268 during rest is preserved in the aged dentate gyrus. BMC Neurosci, 14, 100. doi:10.1186/1471-2202-14-100

Gheidi, A., Satvat, E., & Marrone, D. F. (2012). Experience-dependent recruitment of Arc expression in multiple systems during rest. J Neurosci Res, 90(9), 1820–1829. doi:10.1002/jnr.23057

Guzowski, J. F., McNaughton, B. L., Barnes, C. A., & Worley, P. F. (1999). Environment-specific expression of the immediate-early gene Arc in hippocampal neuronal ensembles. Nat Neurosci, 2(12), 1120–1124. doi:10.1038/16046

Guzowski, J. F., McNaughton, B. L., Barnes, C. A., & Worley, P. F. (2001). Imaging neural activity with temporal and cellular resolution using FISH. Curr Opin Neurobiol, 11(5), 579–584.

Guzowski, J. F., & Worley, P. F. (2001). Cellular compartment analysis of temporal activity by fluorescence in situ hybridization (catFISH). Curr Protoc Neurosci, Chapter 1, Unit 1 8. doi:10.1002/0471142301.ns0108s15

Herdegen, T., & Leah, J. D. (1998). Inducible and constitutive transcription factors in the mammalian nervous system: control of gene expression by Jun, Fos and Krox, and CREB/ATF proteins. Brain Res Brain Res Rev, 28(3), 370–490.

Hunt, S. P., Pini, A., & Evan, G. (1987). Induction of c-fos-like protein in spinal cord neurons following sensory stimulation. Nature, 328(6131), 632–634. doi:10.1038/328632a0

Kron, M. M., Zhang, H., & Parent, J. M. (2010). The developmental stage of dentate granule cells dictates their contribution to seizure-induced plasticity. J Neurosci, 30(6), 2051–2059. doi:10.1523/JNEUROSCI.5655-09.2010

Marrone, D. F., Schaner, M. J., McNaughton, B. L., Worley, P. F., & Barnes, C. A. (2008). Immediate-early gene expression at rest recapitulates recent experience. J Neurosci, 28(5), 1030–1033. doi:10.1523/JNEUROSCI.4235-07.2008

Masuda, N., Ohnishi, T., Kawamoto, S., Monden, M., & Okubo, K. (1999). Analysis of chemical modification of RNA from formalin-fixed samples and optimization of molecular biology applications for such samples. Nucleic Acids Res, 27(22), 4436–4443.

Meltzer, J. C., Grimm, P. C., Greenberg, A. H., & Nance, D. M. (1997). Enhanced immunohistochemical detection of autonomic nerve fibers, cytokines and inducible nitric oxide synthase by light and fluorescent microscopy in rat spleen. J Histochem Cytochem, 45(4), 599–610. doi:10.1177/002215549704500412

Morgan, J. I., & Curran, T. (1989). Stimulus-transcription coupling in neurons: role of cellular immediate-early genes. Trends Neurosci, 12(11), 459–462.

Wilson, M. A., & McNaughton, B. L. (1994). Reactivation of hippocampal ensemble memories during sleep. Science, 265(5172), 676–679.

