## Supplemental Content for "Cellular compartment analysis of temporal activity by fluorescent in *situ* hybridization (catFISH) in the transcardially perfused rat brain"

Image J Fiji Program Cell Counting Protocol

- Open image in program (File -> Open OR Drag and drop from downloads)
- Plugins -> Analyze -> Cell Counter -> Cell Counter -> Initialize -> Type 7
- Begin counting cells stained for positive activity
  - Anything that is bright green in the middle/near-middle of the cell
- Save summary of positive counts:
  - Results (on cell counter tab)
  - File -> Save as
- Image -> Colour -> Split Channels
- Image ->Adjust -> Threshold
  - Keep maximum threshold (bottom bar) at 255
  - Adjust specific threshold (top bar) until cells are visible and not touching
    - Usually, specific threshold value is mid to upper twenties.
- Apply -> Close
- Process -> Binary -> Watershed
- Analyze -> Analyze particles
  - Size: 0.1 – 5
  - Show: Outlines
  - Circularity: 0 – 1.00
- Save summary of negative counts.
